# The *Streptococcus pyogenes* mannose phosphotransferase system (Man-PTS) influences antimicrobial activity and niche-specific nasopharyngeal infection

**DOI:** 10.1101/2024.10.21.619399

**Authors:** Amanda C. Marple, Blake A. Shannon, Aanchal Rishi, Lana Estafanos, Brent D. Armstrong, Veronica Guariglia Oropeza, Stephen W. Tuffs, John K. McCormick

## Abstract

*Streptococcus pyogenes* is a human-adapted pathogen that causes a variety of infections including pharyngitis and skin infections, and although this bacterium produces many virulence and host colonization factors, how *S. pyogenes* competes with the host microbiota is not well understood. Here we detected antimicrobial activity produced from *S. pyogenes* MGAS8232 that was able to prevent the growth of *Micrococcus luteus*. This activity was produced when cells were grown in 5% CO_2_ and in M17 media supplemented with galactose; however, evaluation of the phenotype with the addition of alternative sugars coupled with genome sequencing experiments revealed the antimicrobial phenotype was not related to classical bacteriocins. To further determine genes involved in the production of this activity, a transposon mutant library in *S. pyogenes* MGAS8232 was generated. The transposon screen identified the mannose phosphotransferase system (Man-PTS), a major sugar transporter in *S. pyogenes*, as important for the antimicrobial phenotype. Additional loss-of-function transposon mutants linked to the antimicrobial activity were identified to also be involved in alternative sugar utilization and additionally, the Man-PTS was also further identified from a secondary mutation in a bacteriocin operon mutant. Sugar utilization profiles in all the Man-PTS mutants demonstrated that galactose, mannose, and N-acetylglucosamine utilization was impaired in different Man-PTS mutants. *In vitro* RNA-seq experiments in high and low glucose concentrations further identified the Man-PTS as a glucose transporter; however, there was no transcriptional regulators or virulence factors affected with the loss of the Man-PTS. A clean deletion in the Man-PTS demonstrated defects in a mouse model of nasopharyngeal infection. Overall, the ability of *S. pyogenes* to utilize alternative sugars presented by glycans seems to play a role in acute infection and interactions with the endogenous microbial population existing in the nasopharynx.

**IMPORTANCE:** *Streptococcus pyogenes* causes a wide range of infections and is responsible for over 500,000 deaths per year due to invasive infections and post-infection sequelae. The most common clinical manifestation of *S. pyogenes* however are acute infections such as pharyngitis or impetigo. *S. pyogenes* can adapt to its environment through alternative sugar metabolism and in this study, we identified an antimicrobial phenotype that was not bacteriocin-related but a by-product of alternative sugar metabolism. Evidently, the mannose phosphotransferase system, a well-studied sugar transporter, was involved in production of the antimicrobial, and was also important for *S. pyogenes* to utilize alternative sugars and establish nasopharyngeal infection, but not skin infection. Overall, this study identified potential strategies used by *S. pyogenes* for interactions with the endogenous microbiota and further elucidated the importance of sugar metabolism in acute infection.

---

*Streptococcus pyogenes* (also referred to as group A *Streptococcus*) is a human-adapted bacterial pathogen that commonly colonizes the oropharynx, as well as the skin (1) and it has been found that 5-12% of school-aged children were asymptomatically colonized by *S. pyogenes* (2). *S. pyogenes* also causes a range of acute infections including pharyngitis and impetigo, as well as much more serious invasive infections such as necrotizing fasciitis and streptococcal toxic shock syndrome. Additionally, if repeated infections occur, this can result in autoimmune acute rheumatic fever and rheumatic heart disease (3). It has been estimated that 4 – 8% of the world’s population is affected by pharyngitis each year, and over 500,000 annual deaths globally have been attributed to *S. pyogenes* (4, 5).

Although the pathogenic mechanisms of *S. pyogenes* are well-studied (3), there remains a knowledge gap between the interactions of *S. pyogenes* and the endogenous host microbiota. Bacterial-bacterial competition is often mediated by ribosomally synthesized antimicrobial peptides called bacteriocins that typically exhibit a narrow activity spectrum. *S. pyogenes* can encode an array of bacteriocins including Class I ‘lantibiotic’ bacteriocins such as streptin (6) and salivaricin (7), the non-lantibiotic Class IIb bacteriocins (8–10), and Class III bacteriocin which are specific to M57 serotypes (11). The Class IIb bacteriocin *Streptococcus pyogenes* bacteriocin M (SpbM) is induced during experimental nasopharyngeal infection in mice, suggesting the microenvironment could play an important role in the induction of these bacteriocins (8).

*S. pyogenes* tends to colonize microenvironments that are low in glucose, which includes the nasopharynx and the skin (12–15), and sugar metabolism and transport in *S. pyogenes* has been shown to affect both virulence and host evasion strategies (16–18). *S. pyogenes* carries a collection of complex or alternative sugar transporters and utilization enzymes with ∼15% of the genome being dedicated to sugar metabolism and transport (19–21). These genes encode the enzymes and transporters involved in the phosphoenolpyruvate (PEP)-dependent phosphotransferase systems (PTS) (22). Three enzyme components in these systems undergo phospho-relay events which allow the transport and utilization of an array of carbohydrates (23). The enzyme II (EII) component of the PEP-dependent PTS consists of the PTS transporters that allow bacteria to transport and utilize glucose and other alternative sugars. PTS also encode a cytosolic protein, EIIAB, and a transporter, EIIC. Specifically, the mannose phosphotransferases (Man-PTS) also encodes an additional transport protein, EIID, and the Man-PTS is important in carbohydrate metabolism, virulence regulation, biofilm formation, and immune evasion (17, 24–31). *S. pyogenes* carries 3 mannose phosphotransferases; however, only one of the systems, *manLMN*, is essential for glucose and alternative sugar metabolism, in soft tissue infection and virulence regulation (17, 19, 30).

In this study, we further identify a role for the Man-PTS in colonization and infection in the M18 serotype *S. pyogenes* MGAS8232. Using a transposon mutagenesis library, alternative sugar metabolism was identified to promote the induction of an antimicrobial phenotype, but this was not related to bacteriocin production. While the Man-PTS in *S. pyogenes* did not directly affect virulence or regulation high or low glucose *in vitro* environments, this system was important for *S. pyogenes* to utilize alternative sugars and we show evidence that the Man-PTS aids in niche-specificity during infection as it only affected acute infection in the nasopharynx, but not the skin.

## RESULTS

### *S. pyogenes* produces an antimicrobial compound that is not bacteriocin-related

*S. pyogenes* can encode multiple bacteriocin genes (6–11) and to study bacteriocin induction, we used *S. pyogenes* MGAS8232, an M18 serotype originally isolated from a patient with acute rheumatic fever (32). Based on genomic analysis with BAGEL4 (http://bagel4.molgenrug.nl/), MGAS8232 encodes 3 potential bacteriocins including the Class I bacteriocin salivaricin, although MGAS8232 has a deletion in the *salMT* genes involved in salivaricin production (7, 33). Additionally, two Class IIb bacteriocins (*<u>S</u>treptococcus <u>p</u>yogenes* <u>b</u>acteriocins (*spb*) *JK* and *MN*) are encoded within the genome of MGAS8232, and using a promoter trap strategy, Armstrong *et al* previously demonstrated that a promoter upstream of *spbMN* was induced during experimental nasopharyngeal infection, but not during in vitro growth (8). Therefore, we sought to evaluate environmental cues that may be involved in bacteriocin induction.

To begin to assess different environmental cues that may induce antimicrobial activity, we initially used a set of 9 well-studied bacteriocin indicator strains, including *Micrococcus luteus* (7, 33). Antimicrobial activity from *S. pyogenes* MGAS8232 was determined using the deferred bioactivity antagonism assay. To begin to assess different environmental cues that may induce antimicrobial activity, *S. pyogenes* was grown in Todd Hewitt Broth with 1% yeast (THY) which was considered as a high glucose media, or M17 as a low glucose media. Carbon dioxide (CO_2_) was also examined as an environmental cue that could be a factor in bacteriocin induction as *S. pyogenes* was previously shown to cause differential M-protein transcription in the presence of elevated CO_2_ levels, and is known to be elevated in the upper respiratory system (34, 35). Using the deferred bioactivity antagonism assay, there was no activity when MGAS8232 was grown on standard THY agar in atmospheric or elevated CO_2_ levels (5%), or when grown on M17 agar. However, when M17 media was supplemented with galactose (0.5% (w/v)) as an alternative sugar and grown in elevated levels of CO_2_, inhibition against *M. luteus* was detectable (**Fig. 1A**).

**FIG 1.**
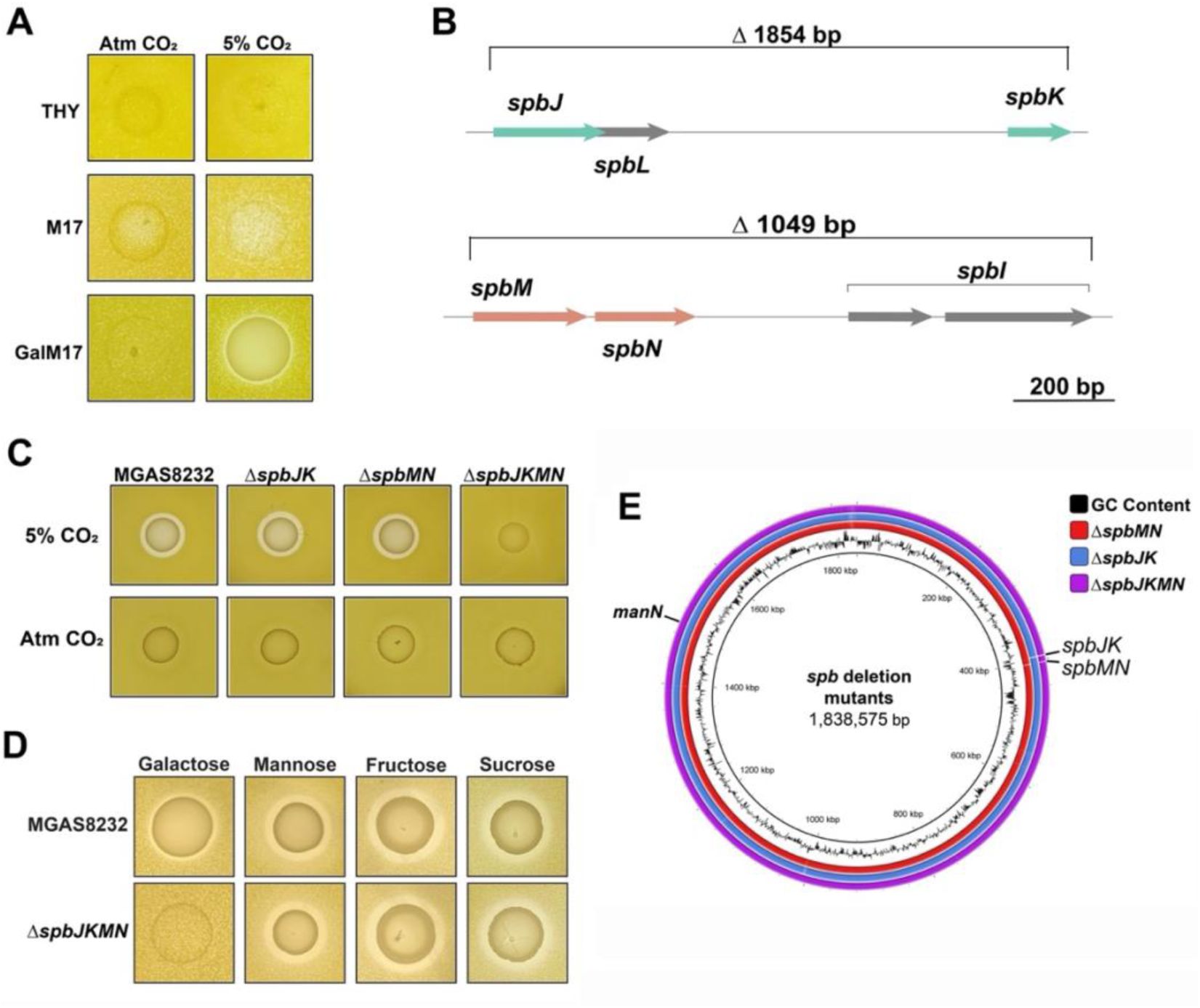
*S. pyogenes* produces an antimicrobial compound that is not related to Class IIb bacteriocins in the presence of alternative sugars and elevated CO_2_. To evaluate antimicrobial activity, the deferred bioactivity antagonism assay was used with the indicator strain *M. luteus*. (A) *S. pyogenes* MGAS8232 was grown on THY, M17 and M17 supplemented with 0.5% galactose and 0.1% CaCO_3_ and grown in either atmospheric conditions (Atm) or 5% CO_2_. (B) Gene schematics of the Class IIb bacteriocin deletions including Δ*spbJK*, Δ*spbMN* and the double deletion strain Δ*spbJKMN*. (C) *S. pyogenes* MGAS8232 and the bacteriocin in-frame deletions were grown on galM17 with 0.1% CaCO_3_ in either Atm CO_2_ or 5% CO_2_. (D) *S. pyogenes* MGAS8232 and the Δ*spbJKMN* in-frame deletion mutant were grown on M17 agar with 0.1% CaCO_3_ containing 0.5% galactose, mannose, fructose or sucrose. (E) Whole genome schematic generated with BRIG of the bacteriocin mutants Δ*spbJK*, Δ*spbMN*, and Δ*spbJKMN* with the *manN* secondary mutation labeled in the Δ*spbJKMN* strain.

As the salivaricin operon contained a *salMT* deletion and is therefore not likely produced in MGAS8232, we generated clean, markerless deletions of either individual Class IIb bacteriocin systems (Δ*spbJK* or Δ*spbMN*), or both bacteriocin systems together (Δ*spbJKMN*), using the *E. coli*/Gram-positive shuttle vector pG^+^host5 system (36) (**Fig. 1B**). Antimicrobial activity was then assessed for the three bacteriocin deletion strains using the deferred bioactivity antagonism assay (**Fig. 1C**). The Δ*spbJK* and Δ*spbMN* deletion strains continued to possess antimicrobial activity. However, the Δ*spbJKMN* strain exhibited a loss in antimicrobial activity (**Fig. 1C**) initially suggesting that both systems were producing this antimicrobial phenotype. However, when investigating other alternative sugars commonly found within the nose via glycans such as mannose and galactose, or dietary sugars introduced in the tonsil environment such as fructose and sucrose, we found that the Δ*spbJKMN* strain was still able to induce an antimicrobial phenotype (**Fig. 1D**). Whole genome sequencing of the Δ*spbJKMN* strain verified the correct deletions, although we also identified secondary mutations in this strain, with one encoding a premature stop codon in the *manN* gene which encodes the transmembrane portion of the mannose phosphotransferase system (Man-PTS) (**Fig. 1E**, **Table S3**). As growth on alternative sugars could still produce the antimicrobial phenotype, and due to the presence of the *manN* mutation, these data suggest the SpbJK and SpbMN peptides were not responsible for this antimicrobial phenotype.

### The antimicrobial phenotype is a product from sugar metabolism

In order to independently identify genes involved in the production of the antimicrobial compound, a transposon library was generated in wildtype *S. pyogenes* MGAS8232 using the <u>K</u>anamycin-<u>r</u>esistant transposon for <u>m</u>assive identification of transposants, Krmit (37). Krmit has been an important genetic tool to investigate essential genes in blood and skin infections of *S. pyogenes* (38, 39). Transposition insertion mutants were screened using the deferred bioactivity antagonism assay against *M. luteus* for a loss of antimicrobial activity (**Fig. 2A**). To determine the insertion site of the transposon in the genome, an arbitrary PCR strategy was used (37), followed by whole genome sequencing to confirm insertion sites and identify any genetic alterations that could have occurred during the transposition event (**Fig. 2B; Table S4**). Interestingly, we identified a loss-of-function mutant that contained a transposon insertion at the beginning of the *manN* gene, similar to the mutation identified in the Δ*spbJKMN* strain. In addition, we identified a transposon insertion in *galC* predicted to encode part of the membrane portion of the galactose PTS as well as *lacA* predicted to encode a galactose-6-phosphate isomerase, involved in the metabolism of galactose and lactose sugars. These findings suggest that the metabolism of alternative sugars was somehow involved in the antimicrobial phenotype against *M. luteus*.

**FIG 2.**
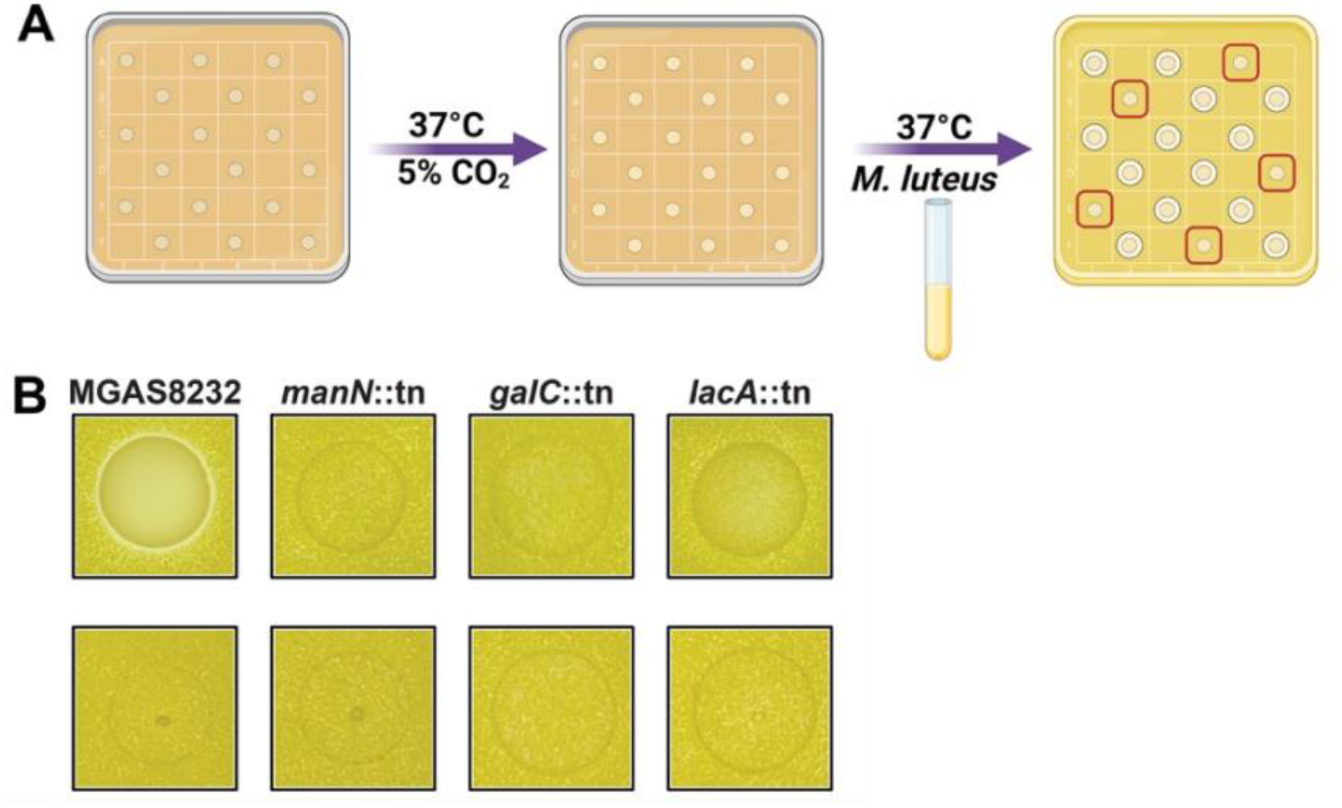
Transposon mutagenesis library screening suggests that sugar metabolism plays an important role in the regulation of the antimicrobial phenotype. (A) Schematic for the *S. pyogenes* MGAS8232 transposon library screening where single colonies were assessed for loss of function using the deferred bioactivity antagonism assay. Isolates were plated on galM17 agar and grown in either elevated CO_2_ or atmospheric CO_2_ levels, and after 24 hours overlayed with soft agar containing *M. luteus*. Red squares highlight no zone of clearance. (B) Deferred antagonism assay with the loss-of-function mutants, with the top row representing the strains that were grown in elevated levels of CO_2_ and the bottom row representing the same strains grown in atmospheric levels of CO_2_.

### Man-PTS is essential for the utilization of alternative sugars by *S. pyogenes*

The Man-PTS has three genes. The *manL* gene is predicted to encode the cytosolic portion of the PTS, while *manM* and *manN* are predicted to encode the transporter. Interestingly, the Man-PTS may also contain a downstream gene annotated as *manO*. In *Streptococcus bovis*, *manO* was believed to be under the control of an independent promoter (40) which may also be present in *S. pyogenes* MGAS8232 (**Fig. 3A**). However, a role for *manO* in *S. pyogenes* is unknown. To investigate if the Man-PTS could have a role in niche-adaptation by *S. pyogenes*, we generated a clean, in-frame deletion of the *manLMN* operon (**Fig. 3A**). The Man-PTS-deficient strain (Δ*manLMN*) was confirmed through PCR (**Fig. 3B**) and whole genome sequencing. Whole genome sequencing did however reveal two mutations resulting in a frameshift in a gene predicted to encode metal ABC transporter ATP-binding protein (SPYM18_RS02175) and a nucleotide change in an intergenic region of the genome (**Table S5**).

**FIG 3.**
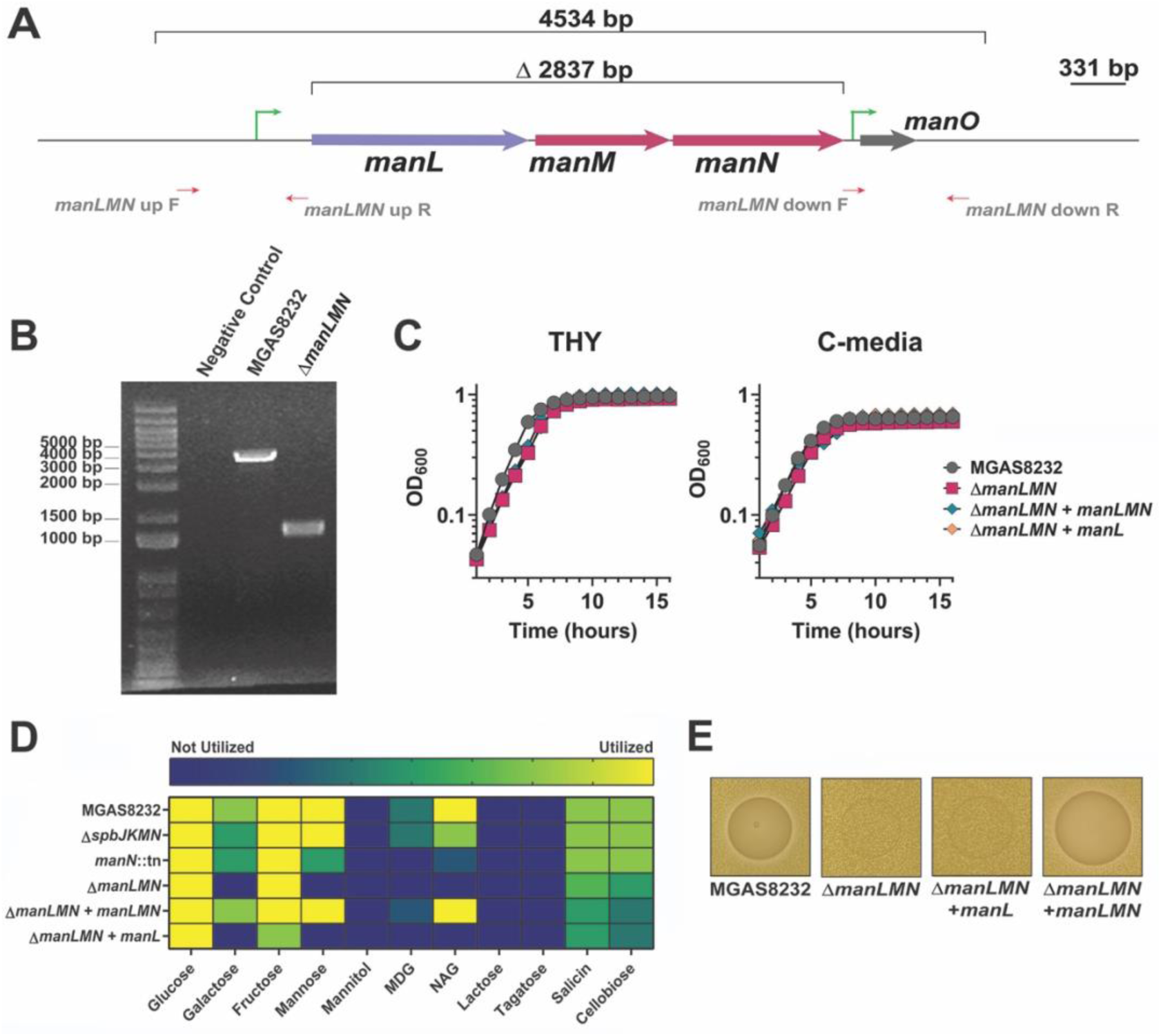
Sugar utilization profile of Man-PTS mutants and genetic complementation in *S. pyogenes* MGAS8232. (**A**) Schematic of the in-frame deletion of the Man-PTS genes (*manLMN*). (**B**) PCR verification of the Man-PTS MGAS8232 Δ*manLMN* deletion strain. (**C**) Growth analyses of wild-type MGAS8232, MGAS8232 Δ*manLMN*, and the genetically complemented strains in both THY and C-media. (**D**) Sugars utilization by wild-type *S. pyogenes* MGAS8232 and the loss of function transposon mutants after 24 hours of incubation at 37°C and 5% CO_2_. Yellow represents the sugar being utilized, green represents partially utilized, blue represents not utilized. MDG-methyl-D-glucopyranoside, NAG – N-acetylglucosamine. (**E**) Deferred bioactivity antagonism assay of wild-type *S. pyogenes* MGAS8232, MGAS8232 Δ*manLMN*, and the genetically complemented strains.

Given these mutations, we genetically complemented the *manLMN* operon using the pDCerm plasmid (41) and cloned either all three genes of the Man-PTS (Δ*manLMN*+*manLMN*) or only the cytosolic portion (Δ*manLMN*+*manL*). Complementation of the cytosolic portion alone was done to determine whether *manL* could restore alternative sugar utilization through other PTSs. This phenomenon has been previously reportered in the M1 serotype of *S. pyogenes*, as different components of the Man-PTS exhibited a differential sugar utilization profiles, and this has been further identified in *Streptococcus mutans* (42). We first ensured that the mutant and complementation clones did not exhibit any defects in growth in either high glucose or low glucose using THY or C-media, respectively (**Fig. 3C**). C-media is a low glucose (0.05%(w/v)) and high peptide concentration media designed to simulate the environment of a deep tissue infection (43). When comparing the Man-PTS mutants Δ*spbJKMN*, *manN*::tn, and the complemented Δ*manLMN* strains, there was notable decreases in the capability to utilize the carbohydrates galactose, mannose and N-acetylglucosamine (**Fig. 3D**). Importantly, the complementation of *manLMN* restored the mutant strains capability to utilize galactose, mannose and N-acetylglucosamine (**Fig. 3D**). However, the complementation of the *manL* did not recover the utilization of these sugars and had a reduction in fructose utilization, suggesting that the Man-PTS needs all three genes to transport these carbohydrates (**Fig. 3D**). Lastly, the Man-PTS deletion strain was unable to produce the antimicrobial when the alternative sugar galactose was present but when all three of the genes were complemented the antimicrobial phenotype was restored, indicating Man-PTS is important for production of the antimicrobial (**Fig. 3E**). Overall, these data indicate that Man-PTS is important in the utilization of alternative carbohydrates, and that all three genes are required to allow these sugars to be utilized and to produce the antimicrobial.

### Man-PTS is important in recognizing glucose in the environment

As Man-PTS has been shown to be important in alternative sugar uptake by *S. pyogenes* (17), we investigated whether the Man-PTS directly or indirectly affected other PTSs and permeases encoded in *S. pyogenes*. To assess this, *S. pyogenes* MGAS8232 and the Δ*manLMN* deletion strain were grown in both THY and C media until they reached stationary growth phase (OD_600_ 0.8) at which point we conducted RNA-seq from both environments (**Fig. S1, Supplemental Table 4&5**). Interestingly, in high glucose, the deficiency of the Man-PTS caused ∼22% of genes involved with PTSs to be significantly upregulated (**Fig. 4A**). These genes are related to PTS components for ascorbic acid, cellobiose, and mannitol; however, the trehalose and fructose PTS were downregulated within the same high glucose environment (**Fig. 4A**). Intriguingly, the fructose-PTS (*fruA*) has been previously shown to not affect fructose utilization or overall virulence (44), while the trehalose PTS was discovered to be involved in the utilization of galactose (17). In the low glucose environment, only the lactose PTS component (EIIA, *lacF.2*) was significantly upregulated in the Man-PTS deficient strain (**Fig. 4B**). These data suggest that the Man-PTS could also be an important glucose transporter in *S. pyogenes* MGAS8232 and in the high glucose environments, the Man-PTS deficient strain could be accommodating glucose uptake using other PTSs.

**FIG 4.**
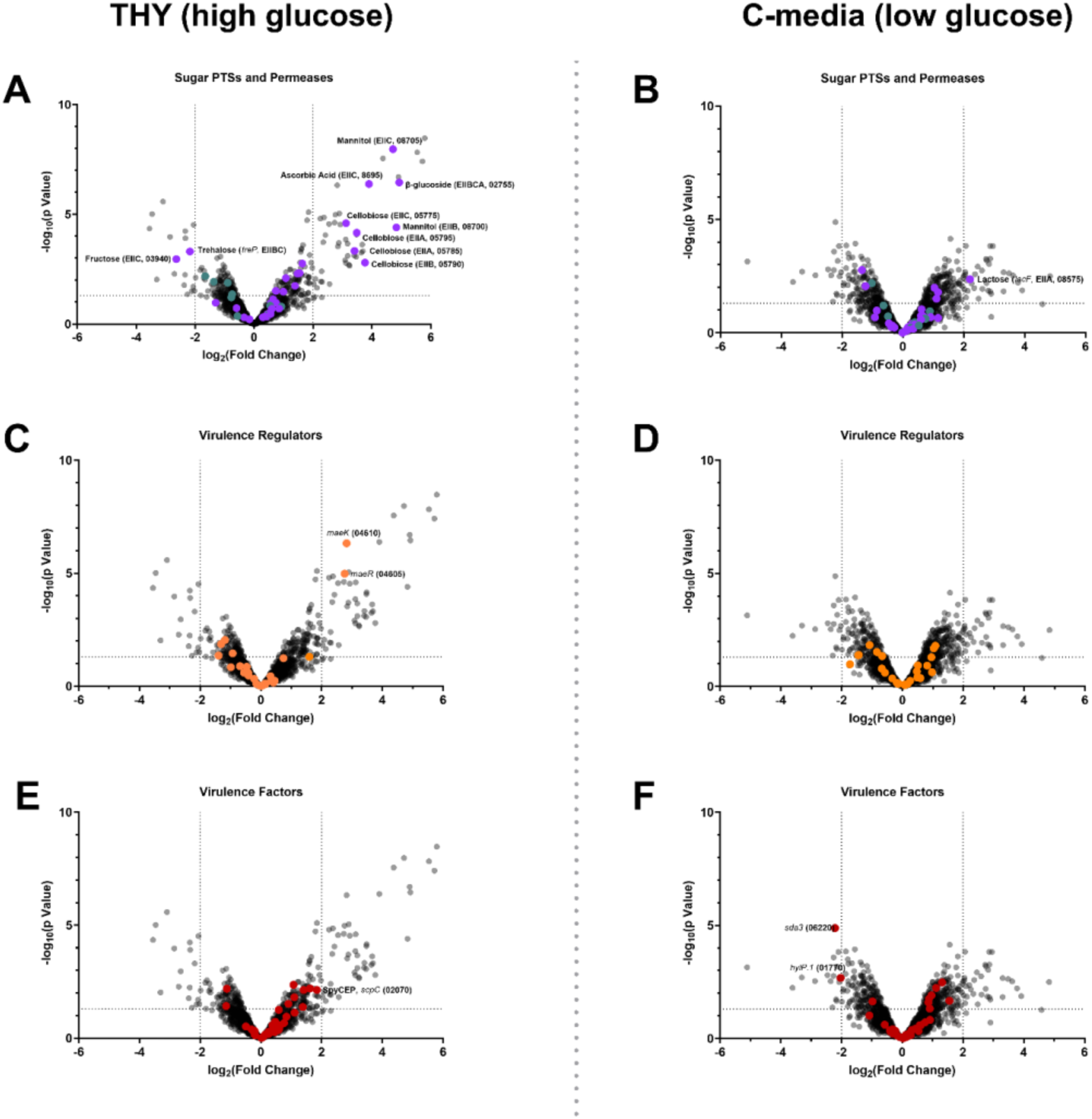
RNA-seq analysis of *S. pyogenes* Δ*manLMN* vs wildtype *S. pyogenes* MGAS8232 in high and low glucose *in vitro* environments. *S. pyogenes* MGAS8232 and Δ*manLMN* were grown in THY (n=3; OD_600_ of 0.7 - 0.8) or C-media (n=3; OD_600_ of 0.6-0.7). All transcripts were compared to the MGAS8232 genome, and each point represents a singular transcript. Genes with a positive log_2_(Fold Change) were increased in the Δ*manLMN* mutant over wild-type *S. pyogenes* MGAS8232. Plots represent labelled transcripts encoding PTSs (purple) and permeases (teal) (**A and B**), two-component system regulators (orange) (**C and D**), and virulence factors (red) (**E and F**). Points above the dottle line represent genes that had significant changes in transcripts between the Δ*manLMN* mutant and wild-type MGAS8232. Quantitation of counts per minute (cpm) are shown in **Fig. S1**.

### Man-PTS does not affect major regulators or virulence factors in *S. pyogenes* MGAS8232

Due to lack of nutrients and multiple stressors from the host, *S. pyogenes* has encoded a variety of regulators that aid in toxin production and immune evasion strategies for niche adaptation, colonization and infection. Previous research has shown that the Man-PTS affected major regulators including carbon catabolite protein A (CcpA) (18, 45), *mga* and *rgg2/3* (30, 46). From the RNA-seq experiment, we investigated transcriptional changes of the two component systems and some major stand-alone response regulators. Many of the regulators were expressed (**Fig. S1**); however, in both *in vitro* environments the loss of the Man-PTS did not cause transcriptional changes in *ccpA*, *mga*, or *rgg2/3*. Furthermore, in high glucose, the loss of the Man-PTS only caused differential transcription with the malate two-component system (*maeRK*) being significantly upregulated (**Fig. 4C**). *S. pyogenes* encodes enzymes involved in the malic enzyme pathway, which permits the utilization of this dicarboxylic acid as an alternative energy source in a CcpA-independent and pH-dependent manner (47). Furthermore, the HPr kinase from the PEP-dependent PTS phosphorylation cascade further promotes the expression malic enzyme pathway (47). Conversely, in the low glucose environment, no virulence regulators exhibited transcriptional changes (**Fig. 4D**). Overall, this data suggests that in high glucose environments and in the absence of Man-PTS, *S. pyogenes* switches to malate as an energy source due to it being unable to adequately import glucose.

*S. pyogenes* carries an arsenal of virulence factors that are uniquely expressed amongst strains. To further investigate whether the Man-PTS affected overall virulence of *S. pyogenes* MGAS8232, we also investigated the transcriptome profiles of key virulence factors in both high and low glucose *in vitro* environments. Firstly, using the Virulence Factor Database (48), 36 virulence factors in *S. pyogenes* MGAS8232 were annotated. In the high glucose environment, there were no virulence factors that were significantly affected in the Δ*manLMN* strain (**Fig. 4E**). However, the *S. pyogenes* cell envelope protease (SpyCEP) was trending to be upregulated within our Man-PTS mutant (**Fig. 4E**, **Fig. S1**). The SpyCEP virulence toxin is known to cleave IL-8 in humans, however it also has the capability to cleave human CXC chemokines, such as CXCL-1/GCP-2, CXCL-6/GROα and mouse CXC chemokines such as CXCL-1/KC and CXCL-2/MIP-2 (49–51). Additionally, there was up-regulation in genes encoding for capsule degradation and decreases in the *hasA* gene, which is involved in capsule production (**Fig. S1**). Previous research has shown that the absence of the capsule in MGAS8232 increases capability of invasion in Detroit-562 pharyngeal cells (52). However, when we infected these cells via a modified gentamycin assay, the Δ*manLMN* strain did not show changes in invasion (**Fig. S2**). Alternatively, in low glucose there was significant down regulation of phage-encoded streptodornase spd3, and hyaluronidase (**Fig. 4F**). Streptodornases allow *S. pyogenes* to evade neutrophils through degradation of neutrophil extracellular traps (NETs), although the *spd3* alone in *S. pyogenes* MGAS5005 was not able to promote skin infection (53, 54) and expression was not completely lost in the Man-PTS deficient strain (**Fig. S1**). Furthermore, the capsule was found to show slight upregulation in the low glucose environment, contrary to the high glucose environment, however this may have been due to relatively low number of transcripts from both wildtype and the Man-PTS deficient *S. pyogenes* MGAS8232. C-media has also been previously shown to induce expression of the cysteine protease, SpeB, however in MGAS8232, *speB* expression was not affected (**Fig. S1**). Overall, the RNA-seq experiments suggested that the Man-PTS does not affect the overall virulence profile of *S. pyogenes* MGAS8232, as many of these virulence regulators and factors are not significantly affected or capable on their own to promote infection.

### Man-PTS is important for nasopharyngeal infection but not for skin infection

One common niche that *S. pyogenes* colonizes is the skin, and previous research has shown that the Man-PTS plays an important role in soft-tissue skin infections (17). To further evaluate this, we compared wildtype *S. pyogenes* MGAS8232 and the Δ*manLMN* mutant in a mouse skin infection model. This model used mice that are transgenic for the expression of major histocompatibility complex (MHC) class II molecules encoding both HLA-DR4 and HLA-DQ8 (herein referred to as B6_HLA_ mice) which sensitizes the mice to the superantigen exotoxins and showed dramatically enhanced nasal infections to *S. pyogenes* MGAS8232 (55). B6_HLA_ mice were subdermally infected with wildtype or the Δ*manLMN* deletion, although no differences in mouse weight, lesion sizes, bacterial burden, or the overall pathology was observed (**Fig. 5**). Overall, these data suggest that the Man-PTS does not play a role in experimental skin infections in *S. pyogenes* MGAS8232.

**FIG 5.**
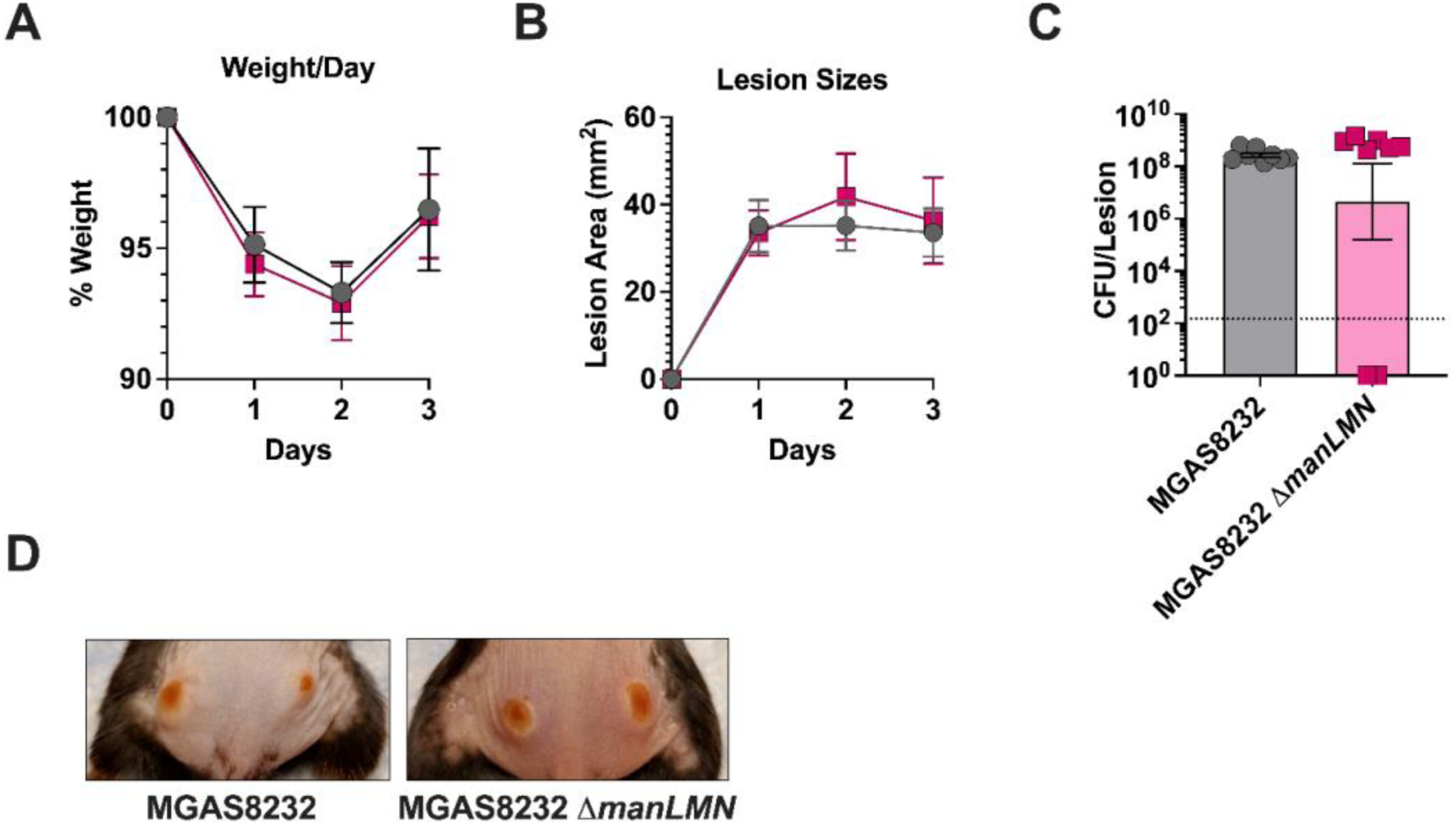
The Man-PTS is not important for establishing skin infection in HLA_B6_ mice. Approximately 5 x 10^7^ CFU of *S. pyogenes* MGAS8232 or Δ*manLMN* were administered intradermally in HLA_B6_ mice (4 mice per group). (**A**) Mouse weights were measured at 24-, 48- and 72-hours post infection and are presented as percentage weight change from day 0. (**B**) Each skin lesion was measured at 24-, 48- and 72-hours post infection. Data points represent lesions in each mouse (4 mice per group, 2 lesions per mouse). (**C**) CFU from each lesion were enumerated at 72 hours post infection and are represented through data points. The dotted line represents the theoretical limit of detection. (**D**) Representative pictures show the respective lesions of the *S. pyogenes* skin infections at 72 hours.

Another primary niche that *S. pyogenes* colonizes is the nasopharynx. To investigate if Man-PTS was important within this niche, B6_HLA_ mice were intranasally infected with either wildtype MGAS8232 or the Man-PTS-deficient Δ*manLMN* strain. We found that at 24- and 48-hours post infection, there was a 1,000- and 10,000-fold reduction in bacterial burden in the complete nasal turbinates, respectively (**Fig. 6**). Although the superantigen streptococcal pyrogenic exotoxin A (SpeA) is known to be essential for nasopharyngeal infection with this strain (55), the gene was not transcriptionally altered in our RNA-seq data in the different glucose concentrations, suggesting the reduction in the bacterial burden was not likely related to SpeA expression (**Fig. S1**). Overall, the Man-PTS in *S. pyogenes* MGAS8232 was important in establishing nasopharyngeal infection in the B6_HLA_ mouse model but was not important for skin infection.

**FIG 6.**
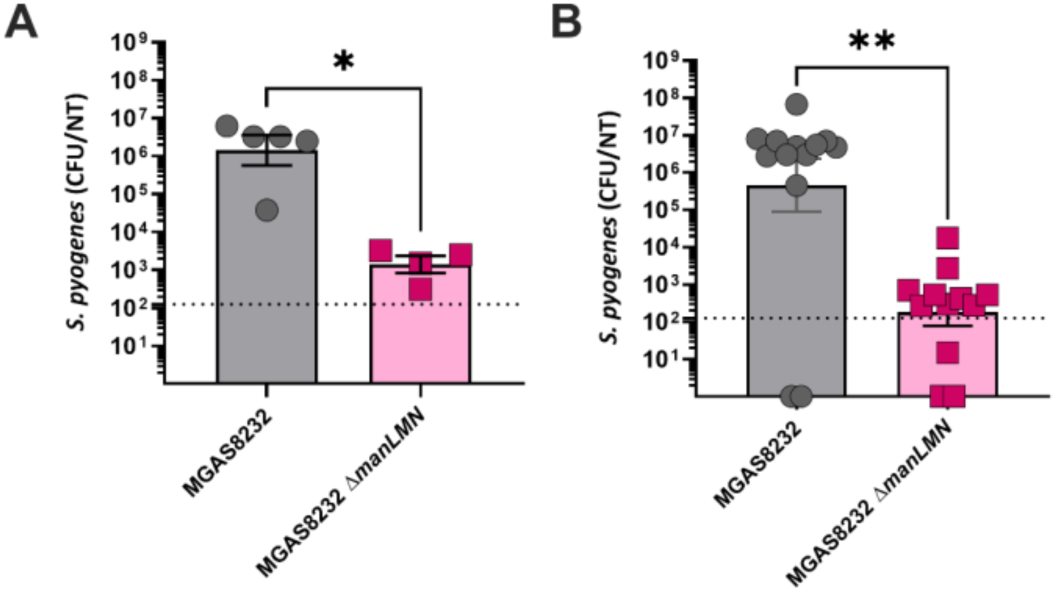
The *S. pyogenes* Man-PTS is important to establish nasal infections at 24- and 48-hours post-infection. HLA_B6_ mice were intranasally inoculated with 1 x 10^8^ CFUs and sacrificed at (**A**) 24- or (**B**) 48 hours post infection with *S. pyogenes* MGAS8232 or the Δ*manLMN* mutant. The bacterial load from the nasal tissues was enumerated and each dot represents an individual mouse. The theoretical yield of detection is illustrated through the horizontal dotted line. * p < 0.01, ** p < 0.001 Mann-Whitney U test.

## DISCUSSION

Commensal and pathogenic bacteria utilize an array of mechanisms to allow for initial survival and coexistence within specific niches (56). *S. pyogenes* is an important human-specific pathogen recognized to often show skin or throat tropism (1). In this study, we provide evidence that alternative sugar metabolism, specifically associated with the Man-PTS system, has a role in establishing acute nasal infection, but not skin infection, providing evidence that sugar metabolism may be important for niche-specificity by *S. pyogenes*.

The nasopharyngeal environment and the surface of skin are known to be depleted for nutrients including glucose, the preferred sugar for *S. pyogenes*. However, alternative sugars are present in these environments (12, 57–59) and *S. pyogenes* can adapt and differentiate its transcriptome in the presence of complex sugars such as fructose, maltose, maltodextrins, and β-glucosides (20, 44, 60, 61). Man-PTS is important for glucose utilization in *S. pneumoniae* (62) and from the RNA-seq experiments, the *S. pyogenes* Man-PTS may play a direct or indirect role in carbon catabolite repression. Interestingly, in high glucose, the Man-PTS deficient strain upregulated the cellobiose PTS, β-glucoside PTS, mannitol PTS, and a component of the ascorbic acid PTS (**Fig. 4A**). In the absence of Man-PTS this may accommodate for inefficient glucose import and its inability to sense its environment. This is further evident in the significant up regulation of the malate two component system in *S. pyogenes* MGAS8232 Δ*manLMN* (**Fig. 4A**), which suggests a switch to malic acid utilization. Interestingly, other than the malate two component system, there was no noted transcriptional differences in other stand-alone or two-component regulators in high or low glucose (**Fig. 4C**).

Interspecies microbe-microbe interactions with *S. pyogenes* and the endogenous microbiota remain understudied and here we discovered antimicrobial activity produced from alternative sugar metabolism by *S. pyogenes*. *S. pyogenes* MGAS8232 encodes Class IIb bacteriocins and the promoter upstream of the bacteriocin gene *spbM* is induced *in vivo* (8) and other research suggests *S. pyogenes* can sense host asparagine to initiate bacteriocin transcription (63). In attempts to characterize additional environmental cues to initiate the production of these bacteriocins, we identified that low glucose and alternative sugar concentrations, and an elevated CO_2_ environment, induced antimicrobial activity from *S. pyogenes* (**Fig. 1**). However, through both unbiased transposon mutagenesis experiments, and then identification of secondary mutations in our double Class IIb bacteriocin deficient Δ*spbJKMN* strain, this phenotype was unrelated to bacteriocin production. The identification of this antimicrobial compound remains unknown.

In other studies, the Man-PTS in the *S. pyogenes* NZ131 strain was found to be involved in the activation and inhibition of quorum-sensing by the regulator Rgg2/3 through the interaction of the stand-alone regulator *mga* via the use of alternative sugars such as mannose and sucrose (30, 46). Our study did not however utilize alternative sugars in the *in vitro* transcriptome analysis and therefore does not account for these potential changes. Importantly, although these *in vitro* transcriptome profiles provide an overall outlook on the role the Man-PTS plays in high and low glucose environments, they do not account for stressors of the immune system or microbiota that would also alter the transcriptome of *S. pyogenes*. Overall, the *in vitro* transcriptome elucidated that the Man-PTS affected overall carbohydrate utilization but did not significantly affect virulence factors or regulators that could affect *S. pyogenes* infection.

Deletion of the Man-PTS system in *S. pyogenes* MGAS5005 caused more aggressive skin infection (17). Using our skin infection model, we did not identify differences in weights, lesion sizes, bacterial burden, or pathology of skin infections in the Man-PTS deficient strain (**Fig. 5**). In terms of virulence affected by the Man-PTS, *S. pyogenes* MGAS8232 Δ*manLMN* demonstrated an increasing trend of SpyCEP transcripts, however skin infection did not display any differences in pathology, as previously shown (49, 51). Previous research has also indicated that the Man-PTS affects the transcription of streptolysin S (17) although this was not observed under our experimental conditions with *S. pyogenes* MGAS8232 (**Fig. S1**). However, in this model *S. pyogenes* was introduced subcutaneously, which is believed to be a high glucose environment and due to this *S. pyogenes* may potentially employ carbon catabolite repression where Man-PTS is not necessary. Furthermore, in a prior *in vivo* transposon mutagenesis study Man-PTS was not essential for soft-tissue invasive infections (38), and in an intact skin infection model, the Man-PTS in *S. pyogenes* NZ131 was not up-regulated but was believed to still play a role in virulence regulation (64). Still, the presence of the Man-PTS is believed to be beneficial to bacteria that colonize mucosal surfaces, like the nasopharynx (65). A pharyngitis infection model in cynomolgus macaques showed upregulation of the *manL* gene at the initial time of acute infection (66). Consistent with this in our mouse nasopharyngeal infection model, the Man-PTS was necessary in establishing acute nasopharyngeal infection at 24- and 48-hours post-infection (**Fig. 6**).

Difference niches colonized by *S. pyogenes* express different glycans on their surface (67, 68). Interestingly, the glycan distribution of the nasopharyngeal tissues comprises of N-glycans which are abundant in galactose, mannose, N-acetylglucosamine and sialic acid (67). Additionally, O-glycans which are abundant in the mucins, only have two carbohydrate differences to them as mannose is absent and fucose is present (68). *S. pyogenes* has been previously theorized to breakdown these glycans to aid in alteration in virulence regulation (46). However, there is currently no evidence of *S. pyogenes* being capable of removing alternative sugars from surface glycans, even though they do encode genes predicted to be glycosyltransferases and glycoside hydrolase proteins that could be used. However, *S. pneumoniae* can do this to cause high concentrations of galactose to be available within the nasopharynx to promote colonization (69). Furthermore, galactose utilization by *S. pyogenes* can aid in the evasion of neutrophil NETs via zinc utilization (16). Man-PTS mutants in *S. pyogenes* MGAS8232 showed deficiencies in the utilization of galactose, mannose, and N-acetylglucosamine (**Fig. 3D**); therefore, as Man-PTS deficient *S. pyogenes* would be unable to sense and thrive using these alternative sugars during the initial steps of nasopharyngeal infection.

This study is the first experimental confirmation that the Man-PTS is required in establishing nasal infection, but not skin infection, suggesting a role in niche specificity. This research also suggests that *S. pyogenes* uses the alternative carbohydrates galactose, mannose, and N-acetyl glucosamine, which are commonly found on glycans of both epithelial cells and mucins, to establish nasopharyngeal infection. This is further evident transcriptionally as the loss of the Man-PTS did not affect the key regulators or the majority of virulence factors in either high or low glucose environments. Research in the exploitation of glycans for carbohydrate utilization should be further explored in the context of *S. pyogenes* infection to identify carbohydrates that could be important in establishing colonization.

## MATERIALS AND METHODS

### Bacterial strains, media, and growth conditions

Bacterial strains and plasmids used in this study are found in **Table 1**. *S. pyogenes* MGAS8232, an M18 serotype isolated from an individual with acute rheumatic fever (32), was grown in Todd Hewitt broth (BD BioSciences) with 1% yeast extract (THY; Thermo Fisher Scientific). When appropriate, 300 µg/mL of kanamycin and/or 100 µg/mL of spectinomycin or 1 µg/mL of erythromycin was added to the media. All molecular cloning was done using *E. coli* XLI-Blue grown in Luria Burtani (LB; Thermo Fisher Scientific) broth or Brain Heart Infusion (BHI; BD Biosciences) supplemented with 50 µg/mL of kanamycin, 100µg of spectinomycin, or 150 µg/mL of erythromycin. All solid media was produced by adding 1.5% (w/v; Thermo Fisher Scientific) of agar.

**Table 1.**
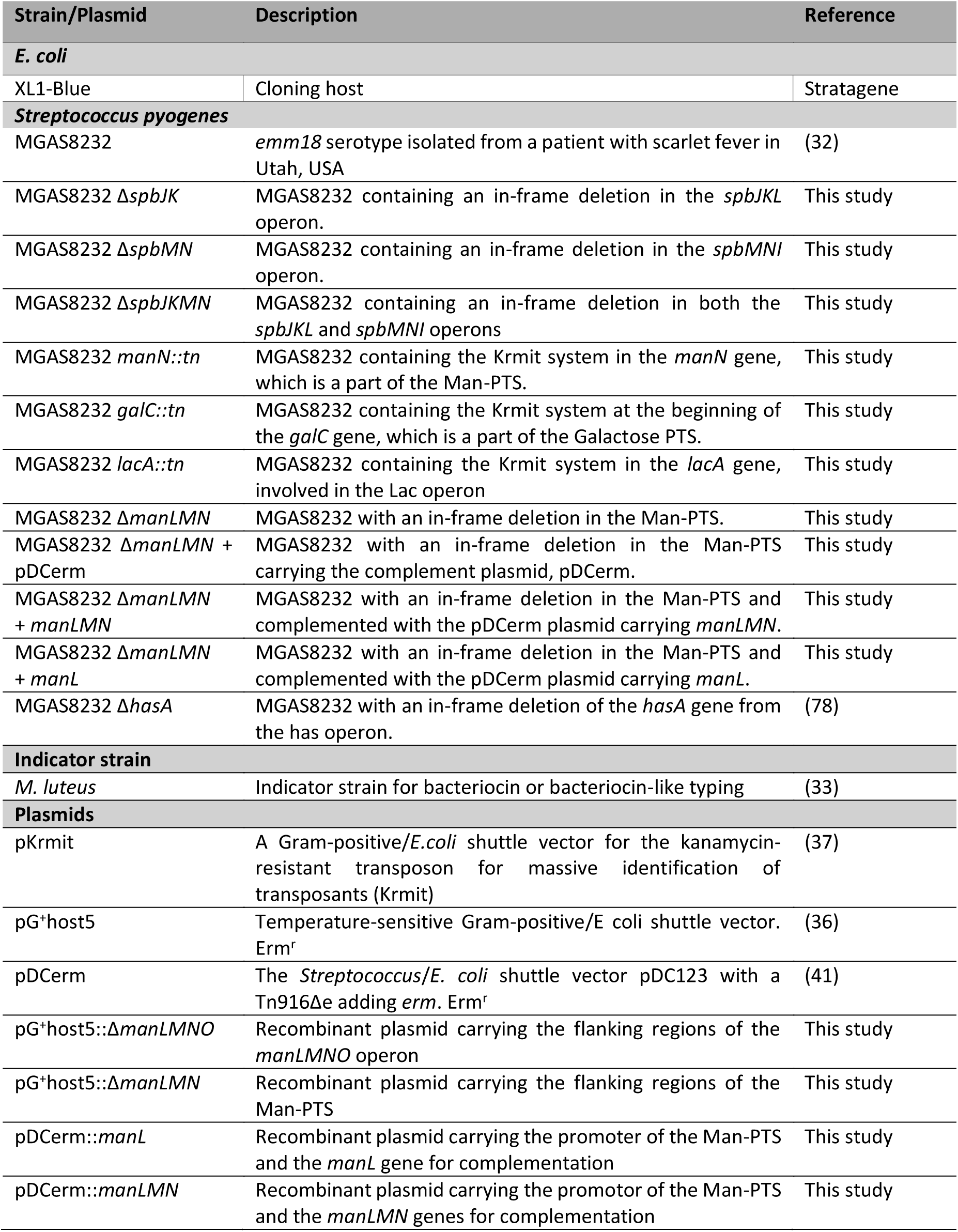
Bacteria and plasmids used in this study.

### Production of in-frame deletion and genetically complemented bacterial strains

To produce deletion mutants in *S. pyogenes*, the Gram-positive *E. coli* temperature-sensitive shuttle vector, pG^+^host5 was used (36). Approximately 500 bp of the upstream and downstream regions (primers are listed in **Table 2**) were cloned into pG^+^host5. The plasmid carrying the flanking regions of the gene of interest was electroporated (Bio-Rad Gene Pulser XCell) into competent *S. pyogenes* and grown at 30°C. Colonies grown on THY with erythromycin (1 µg mL^-1^) at 40°C were selected as single crossover integrations and were further confirmed with PCR. Positive single crossover integrations were grown at 30°C overnight and screened for erythromycin sensitivity. Erythromycin-sensitive colonies are screened with PCR and whole genome sequencing to confirm the deletion and identify any potential mutations that may have occurred during the mutagenesis procedure.

**Table 2.**
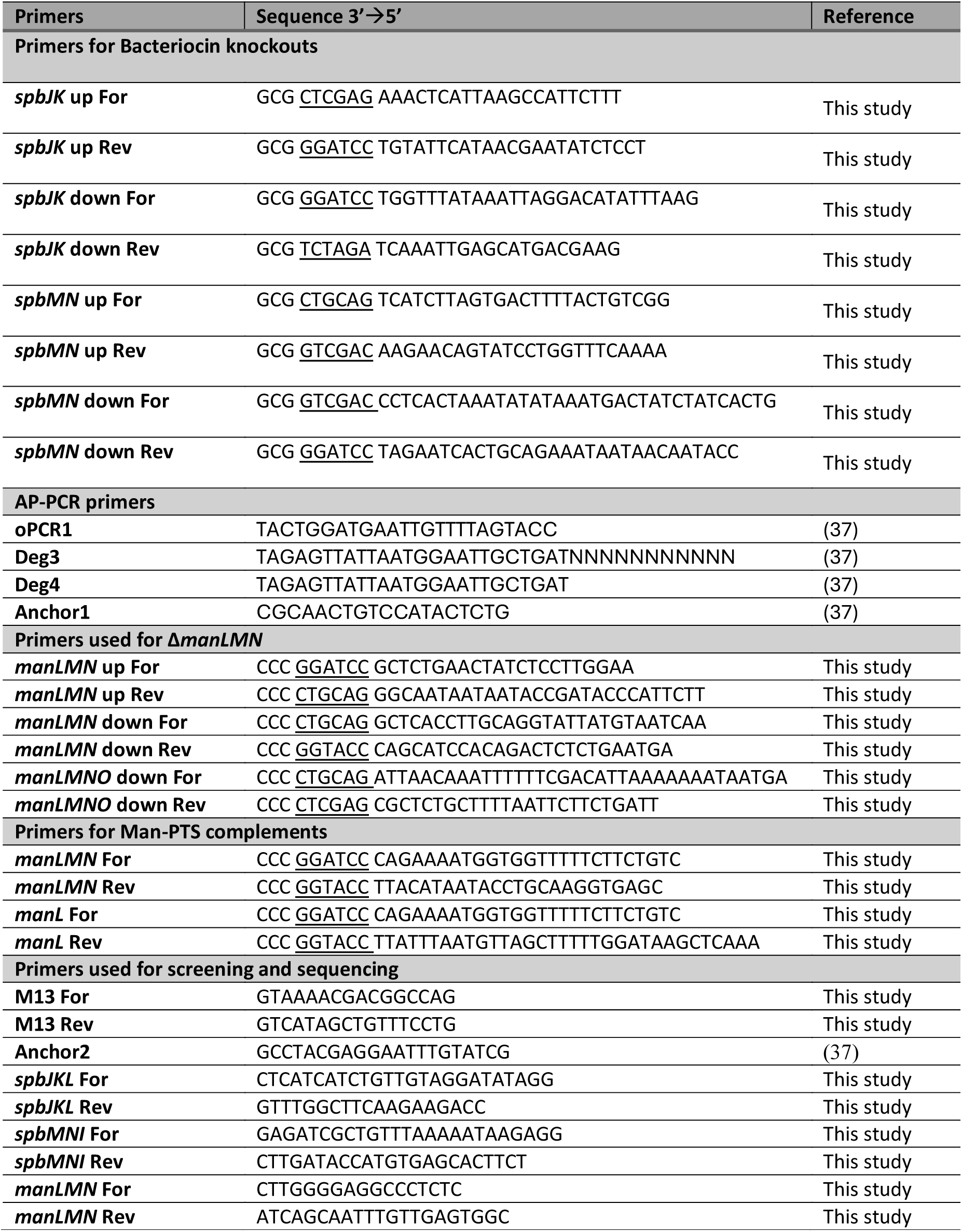
Primers used in this study.

To produce complemented strains with the genes in the Man-PTS, the promoter region was identified using The Berkeley Drosophila Genome Project: Neural Network Promoter Prediction (BDGP: Neural Network Promoter Prediction (fruitfly.org)) and the Softberry: BPROM - Prediction of bacterial promotors (BPROM -Prediction of bacterial promoters (softberry.com)). The promoter with the prospective genes of the Man-PTS were cloned into the complementation plasmid pDCerm [15]. All plasmids were confirmed through DNA sequencing (Plasmidsaurus). The complementation plasmids pDCerm::*manLMN* or pDCerm::*manL* were then electroporated (Bio-Rad Gene Pulser XCell) into electrocompetent *S. pyogenes* MGAS8232 Δ*manLMN* and maintained under antibiotics selection with erythromycin.

### Deferred antagonism bioactivity assay

The deferred antagonism bioactivity assay was used to identify antimicrobial phenotypes from *S. pyogenes* MGAS8232. *S. pyogenes* MGAS8232 was grown on THY or M17 agar (BD Biosciences) supplemented with 0.1% (w/v) calcium carbonate (Thermo Fisher Scientific). M17 plates were supplemented with 0.5% (w/v) alternative sugars as indicated. Calcium carbonate added to the plates ensures that the zones of inhibition were not related to lactic acid production. *S. pyogenes* was grown as previously described and 10 µL of bacterial culture was inoculated on respective M17 agar plates and incubated at 37°C with 5% CO_2_ or atmospheric conditions for 24 hours. The antimicrobial indicator strain, *Micrococcus luteus,* was grown in THY broth and an overnight culture of *Micrococcus luteus* was diluted to an OD_600_ of 1. The diluted culture was then subcultured 1:100 into 42°C THY with 0.7% agar. The inoculated top agar was then overlayed over the bacterial strains of interest and incubated at 37°C overnight. Following overnight incubation, plates were analyzed for a zone of inhibition surrounding MGAS8232, the deletion strains Δ*spbJK*, Δ*spbMN*, Δ*spbJKMN*, and Δ*manLMN*, the transposon mutants, and the complementation strains Δ*manLMN*+*manLMN*, and Δ*manLMN*+*manL*.

### Transposon library production in *S. pyogenes* MGAS8232

To produce a random mutagenesis transposon library in *S. pyogenes* MGAS8232, the pKrmit (37) system was used. The following protocol was adapted and followed from Le Breton *et al* (70). Once the plasmid was electroporated in *S. pyogenes*, transposition frequency was calculated via number of transposition events divided by total number of colonies carrying pKrmit. Non-productive integration was also calculated as the number of spectinomycin resistant transposon mutants over the total number of transposition events. Colonies with a transposition frequency of 10^-2^ – 10^-3^ and a non-productive integration percentage of less than 5% were used. The colonies with the acceptable transposition frequency and non-productive integration percentage were inoculated into 250 mL of THY with 300 µg mL^-1^ of kanamycin and grown for 24 hours at 37°C.

After incubation, this library was stored in THY with 25% glycerol at -70°C. Identification of transposon mutants were conducted through arbitrary primer PCR (71) and whole genome sequencing.

### Whole genome sequencing

1. *S. pyogenes* genomes of the clean deletion mutant, Δ*manLMN* and the transposon mutants sequenced by Seq-Center, LLC. (Pittsburgh, PA), while the bacteriocin clean deletion mutants, Δ*spbJK*, Δs*pbMN*, Δ*spbJKMN*, were sequenced at the London Regional Genomics Centre at the University of Western Ontario for short-read Illumina sequencing. These reads were *de novo* assembled via SPAdes v3.15 (72) and annotated via Prokka v1.12 (73). To determine the presence of single nucleotide mutations, or other mutations, all strains were compared to the reference genome (Genbank accession no. NC_003485) and then compared to the wild-type used in this study using Snippy v4.6.0 (https://github.com/tseemann/snippy).

### *In vitro* RNA isolation and sequencing

To assess the transcriptome between wild-type *S. pyogenes* MGAS8232 and the MGAS8232 Δ*manLMN* strain, RNA was isolated from bacteria grown in high glucose (THY) and low glucose (C-media) media. For the THY media, 6 ml of a *S. pyogenes* overnight culture was added to 100 mL of THY media and the bacteria grew until it reached late exponential phase (OD_600_ of 0.7 – 0.8) at 37°C. For C-media, bacteria were initially grown in THY, washed with PBS and 6 ml was inoculated in 100 ml of C-media and grown late exponential phase (OD_600_ of 0.6 – 0.7). Bacteria were centrifuged at 6000 × *g* for 10 minutes at 4°C, the supernatant was discarded, and the pellet was incubated in RNAProtect Bacteria reagent (Qiagen) following manufacturer’s instructions. Following incubation with RNA-protect, the bacteria were resuspended in 1 mL of 4°C TRIzol (Thermo Fisher Scientific) and stored at -80 °C. TRIzol-treated bacteria were then transferred to tubes containing lysing matrix B beads (MP Biomedicals). Bacteria were homogenized using the Fast Prep-24 Classic grinder and lysing machine (MP Biomedicals) twice at 6.5 m/s for 45 seconds with a 90 second break on ice in-between. After homogenization, 0.2 volumes of chloroform were added, and samples were incubated on ice for 10 minutes. To isolate the RNA from the aqueous layer, the solution was centrifuged at 14,000 × *g* for 15 minutes at 4 °C, the colourless aqueous layer at the top was collected, and an equal amount of 70% ethanol was added. RNA was then isolated through the PureLink RNA mini kit (Thermo Fisher Scientific), and the contaminating DNA was removed with the PureLink DNase set (Thermo Fisher Scientific) following the manufacturer’s instructions. RNA was eluted in 56°C MilliQ water and stored at -80 °C.

An Agilent 2100 Bioanalyzer at London Regional Genomics Centre was used to determine the RNA integrity number (RIN), and samples with an RIN above 8 were sequenced by 12 million paired-end Illumina RNA sequencing (Seq-Center). To map the reads, HISAT2 (74) was used. To normalize counts for each gene and to produce the counts per million (cpm), all reads were inputted to R (https://www.R-project.org/) and edgeR’s (75) Trimmed Mean of M values (TMM) algorithm. The differential expression analysis amongst two groups was done in EdgeR’s glmQLFTest, which provided the subsequent log_2_(fold change) and the p values for each gene based on the cpm. The reference genome used for the comparisons and mapping of the sequencing was the publicly available *S. pyogenes* MGAS8232 genome (Genbank accession no. NC_003485).

### Sugar utilization profiling

To determine the sugar utilization profiles for *S. pyogenes* the API® 50CH system (Biomerieux) was used. This system allows tests for the utilization of 49 different sugars. *S. pyogenes* MGAS8232, the Δ*spbJKMN*, *manN*::*tn*, Δ*manLMN* mutants, and the complemented Man-PTS strains, were grown on THY agar with their respective antibiotic concentrations overnight at 37°C. After incubation, bacteria were scraped from the plates and transferred to Hank’s Balanced Salt Solution (HBSS) to an OD_600_ of approximately 3.0 and then diluted in the CHL medium (Biomerieux) to OD_600_ of 0.14. Next, 200 µL of this solution was inoculated to each cupule of the API® strips. To ensure that *S. pyogenes* was grown in an anaerobic environment, 50 µL of mineral oil was inoculated to the top of each cupule. The API® 50CH system was then protected from the light and incubated for 24 hours at 37 °C with 5% CO_2_. The CHL medium (Biomerieux) contains the pH indicator bromocresol purple, and sugar utilization by the bacteria results in a decrease in pH due to the production of acid.

### Human cell culture and invasion assays

The Detroit-562 human pharyngeal cell line (ATCC CCL-138) was maintained in minimal essential medium eagle (MEM;) at 37°C in 5% CO_2_. The MEM was supplemented with 10% (v/v) heat-inactivated fetal bovine serum (FBS; Sigma-Aldrich), 1 mM of GlutaMAX™ supplement (Thermo Fisher Scientific), and 100 µg mL^-1^ streptomycin and 100 U mL^-1^ penicillin (Life Technologies). The invasion assay is a modified gentamycin protection assay to evaluate the number of bacteria internalized by the Detroit-562 cells. Cells were grown to ∼90% confluence in 12 well tissue culture treated plates (Falcon, Corning). Monolayers were washed three times with PBS and replaced with antibiotic-free MEM. *S. pyogenes* was grown to early exponential phase (OD_600_ = 0.2 – 0.4) and each well was supplemented with bacteria at a multiplicity of infection (MOI) of 100 based on the calculated average number of cells in the wells. Inoculated cell culture plates were incubated for 2.5 hours at 37 °C in 5% CO_2_. After incubation, monolayers were washed, and MEM supplemented with 100 µg mL^-1^ gentamycin (Sigma-Aldrich) was added and incubated for 1.5 hours at the same conditions previously stated to kill remaining extracellular bacteria. After incubation, cells were washed and lysed with 400 µL of cold 0.1% Triton X-100 (VWR International) for 5-10 minutes. To enumerate the number of internalized bacteria, the bacteria were serially diluted and plated on TSA 5% sheep blood.

### Animal ethics statement

Mouse experiments conducted were in accordance with the Canadian Council of Animal Care Guide to the Care and Use of Experimental Animals. The Animal Use Protocol (AUP) number 2020-041 was approved by the Animal Use Subcommittee at the University of Western Ontario (London, ON, Canada).

### *S. pyogenes* skin infection

8-12 week old B6_HLA_ mice were used in the skin infection model. Mice were shaved using clippers and any remaining fur was removed using commercial hair removal cream, 24 hours prior to infection. Mice were subcutaneously inoculated with 100 µL, with 2.5 x 10^7^ CFU of bacteria (50 µL) into each flank. Mice were weighed daily, and the lesion sizes were measured using calipers. Mice were sacrificed at 72 hours post-infection and the lesions were harvested and homogenized. To enumerate the number of bacteria in each lesion, the homogenized lesions were serially diluted and plated on tryptic soy agar (TSA) supplemented with 5% sheep blood.

### Acute nasopharyngeal infection

8-12 week old B6_HLA_ mice were used to establish acute *S. pyogenes* infection within the nasopharynx (55, 76, 77). Bacteria were grown to early exponential phase (OD_600_ 0.2-0.4) and washed with HBSS twice. Each nostril was inoculated with 7.5 µL of bacterial inoculum, for a total of approximately 1-2 x 10^8^ colony forming units (CFUs), under Forane (isoflurane; USP) inhalation anaesthetic (Baxter Corporation, Canada). At 24 or 48 hours post infection, mice were sacrificed, and their nasal tissues were extracted. To enumerate the bacteria within the nasal tissues, tissues were homogenized, serially diluted, and plated on TSA with 5% sheep blood and grown in 37 °C overnight. Counts considered below the limit of detection had less than 30 CFU per 100 µL of nasal tissues.

## Notes

### Competing Interest Statement

The authors have declared no competing interest.

