## Supplementary Figs 1,2 Tables 1, 2 3 for "The *Streptococcus pyogenes* mannose phosphotransferase system (Man-PTS) influences antimicrobial activity and niche-specific nasopharyngeal infection"

Short title: *S. pyogenes* sugar metabolism and niche specificity

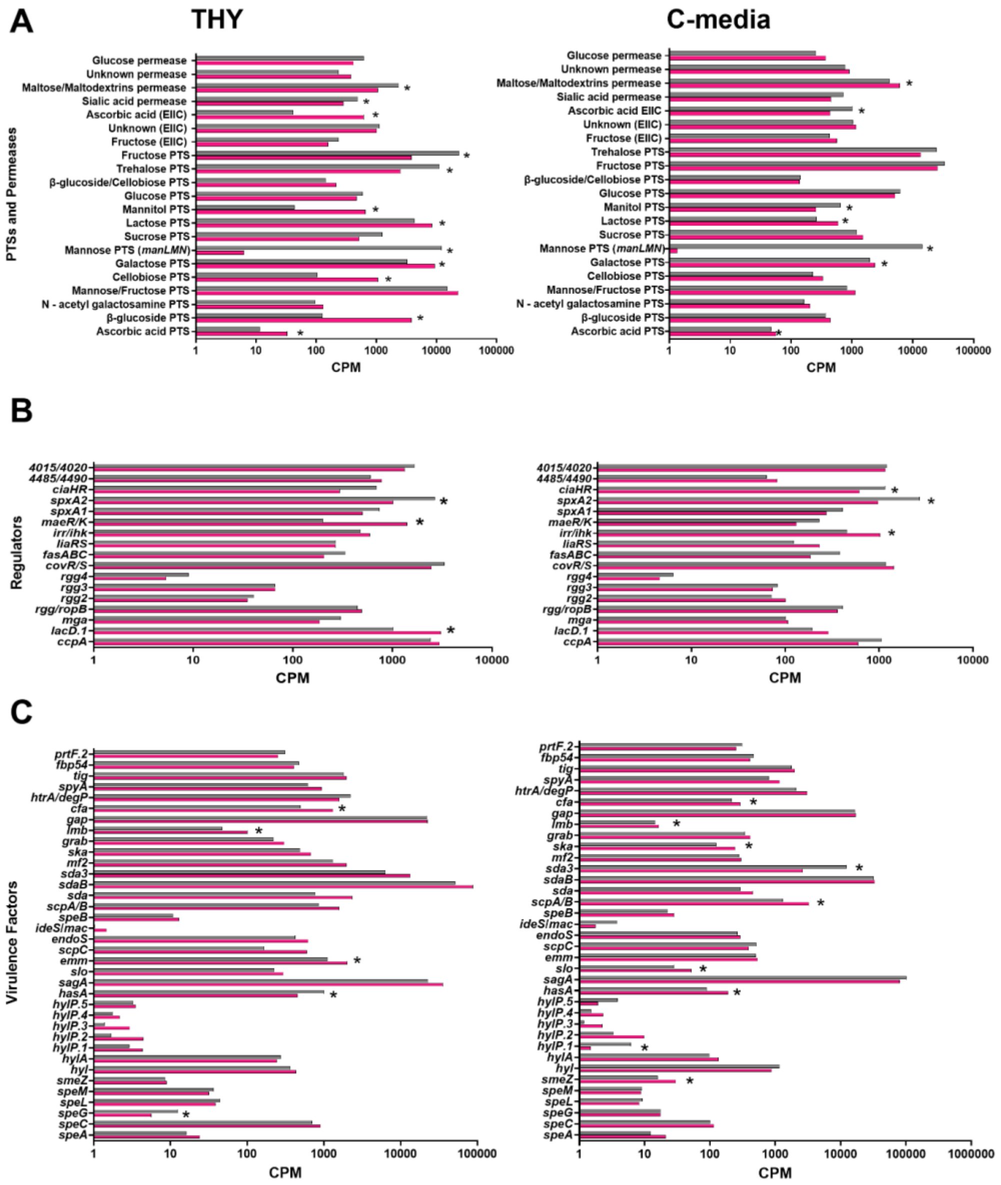


**FIG S1. Transcripts of sugar PTSs and permeases, virulence regulators, and factors in wildtype *S. pyogenes* MGAS8232 and the Δ*manLMN* strain when grown in high and low glucose environments.** *S. pyogenes* MGAS8232 and Δ*manLMN* were grown in THY (n=3; $OD_{600}$ of 0.7 - 0.8) or C-media (n=3; $OD_{600}$ of 0.6-0.7). Samples with RNA integrity numbers (RIN) of ≥ 7 were subjected to RNA-seq. The following represents the transcripts (counts per million (CPM)) from genes related to each (**A**) sugar PTS and permease, (**B**) virulence regulators, and (**C**) virulence factors that are encoded in S. pyogenes MGAS8232. All virulence factors were identified via the virulence factor database (VFDB: Virulence Factors of Bacterial Pathogens (mgc.ac.cn)). PTSs, permeases, and virulence regulators that consist of more than one gene are represented by the sum of their transcripts. The transcripts from wildtype MGAS8232 and the Δ*manLMN* strain are represented in grey and pink bars, respectively. * represents at least one gene that had $p \leq 0.05$.

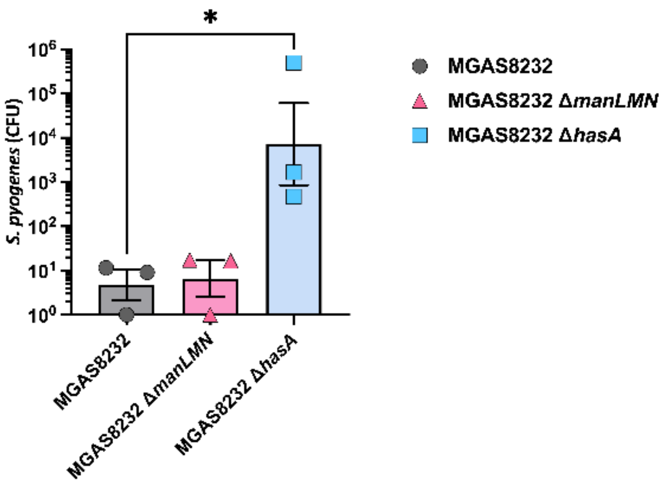


**FIG S2. The reduction of capsule in *S. pyogenes* Δ*manLMN* does not cause increased invasion in Detroit-562 cells.** Internalization assay of wild-type *S. pyogenes* MGAS8232 via a modified classical gentamycin assay comparing the Man-PTS (Δ*manLMN*) and hyaluronic acid capsule (Δ*hasA*) deficient mutants in Detroit 562 cells. D562 cells were cultured with *S. pyogenes* (MOI 100) for 2 hours at 37°C with 5% CO2, then supplemented with 100µg/mL of gentamycin for 1.5 hours. Bars represent geometric mean, and each dot is a biological replicate. *, $p \leq 0.1$ via One-Way ANOVA.

**Supplemental Table 1. Single nucleotide polymorphisms and genetic alterations in the Spb clean deletion mutants**

| | **Single Nucleotide Polymorphisms** | | | | | | | |
|---|---|---|---|---|---|---|---|---|
| | **Position** | **Type** | **Ref** | **Alt** | **AA change** | **Effect** | **Gene** | **Locus tag** |
| ***ΔspbJK*** | 354777 | del | CT | C | Ser27fs | frameshift | HlyC/CorC family transporter | SPYM18_RS01955 |
| | 1873136 | del | AT | A | Ile140fs | frameshift | insulinase family protein | SPYM18_RS09550 |
| ***ΔspbMN*** | 96100 | snp | T | C | Pro619Pro | synonymous | *rpoB*, DNA-directed RNA polymerase subunit beta | SPYM18_RS00630 |
| | 1450278 | snp | T | C | Met410Val | missense | NCS2 family permease | SPYM18_RS07500 |
| | 1753971 | snp | T | C | Glu47Gly | missense | streptopain | SPYM18_RS09005 |
| ***ΔspbJKMN*** | 96100 | snp | T | C | Pro619Pro | synonymous | *rpoB*, DNA-directed RNA polymerase subunit beta | SPYM18_RS00630 |
| | 987920 | snp | G | T | Ala499Glu | missense | ABC transporter ATP-binding protein | SPYM18_RS05100 |
| | 1140999 | snp | G | A | Ala83Val | missense | LPXTG cell wall anchor domain-containing protein | SPYM18_RS05925 |
| | 1421847 | snp | T | C | Thr225Ala | missense | Na/Pi cotransporter family protein | SPYM18_RS07330 |
| | 1450278 | snp | T | C | Met410Val | missense | NCS2 family permease | SPYM18_RS07500 |
| | 1496942 | snp | G | A | Trp117* | stop gained | *manN*, PTS mannose/fructose / sorbose transporter family subunit IID | SPYM18_RS07830 |
| | 1498186 | snp | G | T | N/A | N/A | N/A | N/A |
| | 1753971 | snp | T | C | Glu47Gly | missense | streptopain | SPYM18_RS09005 |
| | 1763350 | snp | G | A | N/A | N/A | N/A | N/A |
| | 1765535 | del | CT | C | N/A | N/A | N/A | N/A |
| | 1873136 | del | AT | A | Ile140fs | frameshift | insulinase family protein | SPYM18_RS09550 |

Note: AA=amino acid, del=deletion, fs=frameshift, snp=single nucleotide polymorphism, *=stop codon.

**Supplemental Table 2. Single nucleotide polymorphisms and genetic alterations in the transposon mutants**

| | Single Nucleotide Polymorphisms | | | | | | | |
|---|---|---|---|---|---|---|---|---|
| | **Position** | **Type** | **Ref** | **Alt** | **AA change** | **Effect** | **Gene** | **Locus tag** |
| ***manN*::tn (07830)** | 229153 | snp | C | T | Ser828Phe | missense | SEC10/PgrA surface exclusion domain-containing protein | SPYM18_RS02175 |
| | 1364445 | snp | G | A | Thr553Ile | missense | primosomal protein N' | SPYM18_RS07080 |
| | 1675147 | snp | G | A | Ala413Val | missense | *pnp*, polyribonucleotide nucleotidyltransferase | SPYM18_RS08685 |
| | 1873457 | del | AT | A | Ile33fs | frameshift | insulinase family protein | SPYM18_RS09550 |
| ***galC*::tn (07380)** | 229153 | snp | C | T | Ser828Phe | missense | SEC10/PgrA surface exclusion domain-containing protein | SPYM18_RS02175 |
| | 1364445 | snp | G | A | Thr553Ile | missense | primosomal protein N' | SPYM18_RS07080 |
| | 1390878 | snp | G | A | Ala134Val | missense | amino acid ABC transporter permease | SPYM18_RS07200 |
| | 1675169 | snp | G | A | His406Tyr | missense | *pnp,* polyribonucleotide nucleotidyltransferase | SPYM18_RS08685 |
| | 1873457 | del | AT | A | Ile33fs | frameshift | insulinase family protein | SPYM18_RS09550 |
| ***lacA*::tn (08595)** | 229153 | snp | C | T | Ser828Phe | missense | SEC10/PgrA surface exclusion domain-containing protein | SPYM18_RS02175 |
| | 1364445 | snp | G | A | Thr553Ile | missense | primosomal protein N' | SPYM18_RS07080 |
| | 1420865 | snp | G | A | N/A | N/A | N/A | N/A |
| | 1873457 | del | AT | A | Ile33fs | frameshift | insulinase family protein | SPYM18_RS09550 |

Note: AA=amino acid, snp=single nucleotide polymorphism, del=deletion, fs=frameshift, N/A=not available.

**Supplemental Table 3 Single nucleotide polymorphisms and genetic alterations in the Man-PTS deficient strain.**

| | Single Nucleotide Polymorphisms | | | | | | | |
|---|---|---|---|---|---|---|---|---|
| | **Position** | **Type** | **Ref** | **Alt** | **AA change** | **Effect** | **Gene** | **Locus tag** |
| ***ΔmanLMN*** | 401924 | del | GA | G | Lys110fs | frameshift | metal ABC transporter ATP-binding protein | SPYM18_RS02175 |
| | 1263863 | snp | G | A | N/A | N/A | N/A | N/A |

Note: AA=amino acid, del=deletion, fs=frameshift, snp=single nucleotide polymorphism.
